# Structure and Mechanism of Avermitilol Synthase, a Sesquiterpene Cyclase that Generates a Highly Strained 6-6-3 Tricyclic Alcohol

**DOI:** 10.1101/2025.09.11.675706

**Authors:** Matthew N. Gaynes, Kristin R. Osika, David W. Christianson

**Author notes:** Corresponding author: David W. Chrisianson.

## Abstract

Avermitilol synthase from *Streptomyces avermitilis* (SaAS) is a high-fidelity class I terpene cyclase that converts farnesyl diphosphate into a highly-strained, 6-6-3 tricyclic sesquiterpene alcohol. The mechanism of avermitilol formation proceeds through a 10-3 bicyclic intermediate, bicyclogermacrene, which undergoes proton-initiated anti-Markovnikov opening of two separate C=C bonds in a transannulation mechanism that forms the 6-6-3 tricyclic skeleton, with quenching by water to yield avermitilol. Small amounts of a side product, viridifloral, result from Markovnikov opening of one of the reactive C=C bonds. Here, we present enzymological studies of SaAS to establish the substrate scope and metal ion dependence for catalysis, and we present crystal structures of SaAS complexed with a variety of ligands that partially mimic carbocation intermediates in catalysis. Interestingly, these structures show that two water molecules remain trapped in a polar crevice in the active site regardless of the ligand bound. Structure-activity relationships for site-specific mutants yield key insight on the catalytic importance of these trapped water molecules. Specifically, T215 normally hydrogen bonds with water molecule W1, but the T215V substitution breaks this hydrogen bond and causes W1 to shift by 1.3 Å to form a hydrogen bond with W300. Avermitilol generation is completely blocked in this mutant, but the generation of viridifloral and another side product is enhanced. Thus, the T215V substitution causes water molecule W1 to align for reaction with the tertiary and not the secondary carbon in the reactive C=C bond of bicyclogermacrene.

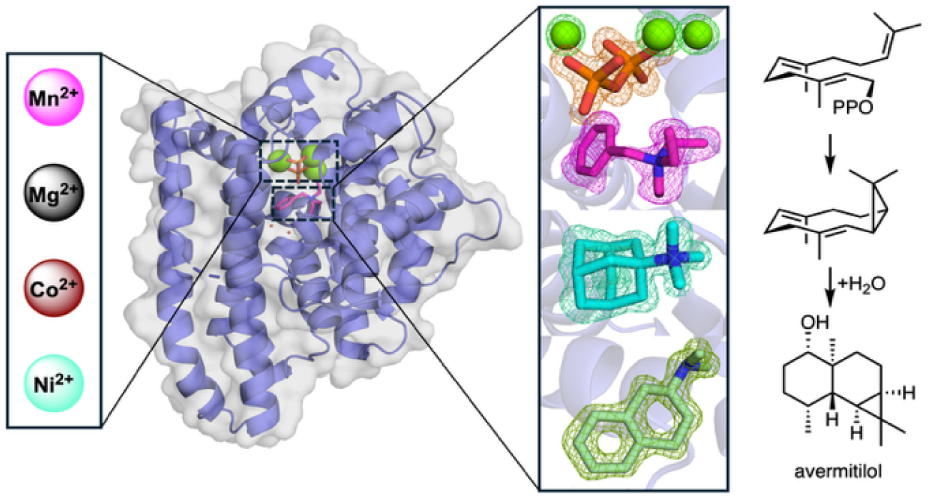

## Introduction

The biosynthesis of terpene natural products is a striking example of nature’s synthetic capability, featuring multi-step enzyme-catalyzed reactions that typically generate complex polycyclic scaffolds with precise regiochemistry and stereochemistry.^1–3^ Many terpene natural products are commercially valuable and have diverse applications. For example, the sesquiterpenes artemisinin, bisabolane, and nootkatone have important applications in pharmaceutical, energy, and environmental sciences, respectively.^3–5^ Sesquiterpenes derive from a linear, achiral C_15_ substrate, farnesyl diphosphate (FPP), which undergoes cyclization in a reaction catalyzed by a sesquiterpene synthase.^7,8^

A class I sesquiterpene synthase has two conserved metal binding motifs at the mouth of its active site that coordinate to a trinuclear metal cluster.^9^ The substrate diphosphate group also coordinates to this metal cluster, which triggers dissociation to yield inorganic pyrophosphate (PP_i_) and an allylic carbocation intermediate. The distinct shape of the active site pocket enforces a unique conformation of the flexible isoprenoid substrate so that the allylic carbocation (usually C1) reacts with one of the remaining π bonds in concert with or subsequent to PP_i_ departure. The C1–C6, C1–C10, or C1–C11 ring closure reactions lead to the bisabolyl cation, germacryl cation, or humulyl cation, respectively.^10–15^ A well-defined sequence of carbon-carbon bond-forming steps follows initial carbocation formation; reaction sequences are ultimately terminated by proton elimination to yield a polycyclic alkene such as *epi*-isozizaene, or by addition of a water molecule to yield an alcohol such as bisabolol.^11,15^

Enzymes that catalyze anti-Markovnikov reactions, i.e., a carbon-carbon bond-forming reaction that yields a less-stable secondary carbocation rather than a more-stable tertiary carbocation, are of particular interest since they defy traditional rules of organic chemistry. While computational studies indicate that relatively few secondary carbocations intermediates exist as energy minima in terpene cyclization reactions, C1–C11 cyclization reactions of farnesyl diphosphate yield a secondary humulyl carbocation that can be stabilized by intramolecular effects such as hyperconjugation.^16^ The prototypical example of such a reaction step is found in the reaction mechanism of pentalenene synthase, the first enzyme identified to catalyze initial C1–C11 cyclization.^12,13,17–20^ Structural and enzymological studies pinpoint key active site interactions and residues that stabilize carbocation intermediates to promote anti-Markovnikov ring closure. Subsequently, structural and enzymological studies of cucumene, caryolan-1-ol, longiborneol, Δ^6^-protoilludene, presilphiperfolan-8β-ol, and trichobrasilenol synthases similarly reveal active site features that appear to facilitate initial anti-Markovnikov C1–C11 cyclization in FPP cyclization cascades.^14,15,21,22^

Anti-Markovnikov reactions can also occur in the middle rather than the beginning of a terpene cyclization sequence. For example, avermitilol synthase from *Streptomyces avermitilis* (SaAS) utilizes FPP to generate the eponymous 6-6-3 tricyclic sesquiterpene alcohol (Figure 1A).^23^ The proposed reaction mechanism of SaAS proceeds through initial C1–C10 ring closure followed by deprotonation at C1 to yield a cyclopropane intermediate, bicyclogermacrene. Anti-Markovnikov opening of one C=C bond in bicyclogermacrene enables transannulation via anti-Markovnikov opening of the second C=C bond; addition of a water molecule quenches the reaction to yield avermitilol.^24^ Minor side products germacrene A and germacrene B result from premature termination of the cyclization cascade by proton elimination, and minor side product viridifloral results from an alternative transannulation reaction (Figure 1A).

**Figure 1.**
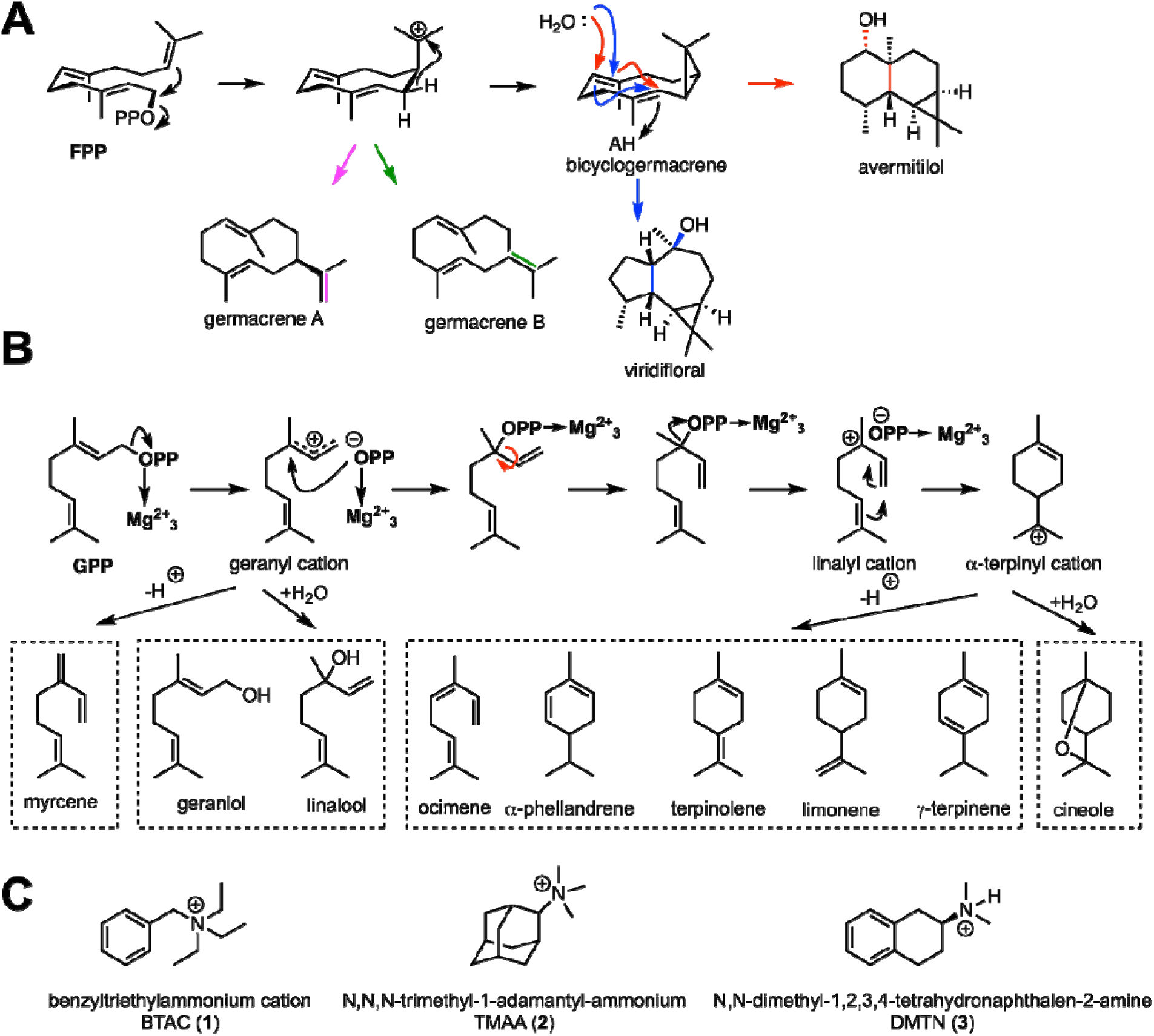
Terpene cyclization pathways catalyzed by SaAS. (A) Native FPP cyclization pathway leading to avermitilol, plus minor derailment reactions leading to viridifloral and germacrenes A and B. Of note, anti-Markovnikov opening of the two C=C bonds of bicyclogermacrene is required for avermitilol generation. (B) Monoterpene reaction pathways yield multiple linear, cyclic, and hydroxylated products. (C) Ligands used for co-crystallization experiments.

We now report the high-resolution X-ray crystal structures of wild-type SaAS in complex with a Mg^2+^_3_-PP_i_ cluster and (1) the benzyltriethylammonium cation (BTAC), (2) *N,N,N*-trimethyl-1-adamantyl-ammonium hydroxide (TMAA), and (3) *N,N*-dimethyl-1,2,3,4-tetrahydronaphthalen-2-amine hydrochloride (DMTN) (Figure 1C), as well as crystal structures of Mg^2+^_3_-PP_i_-BTAC complexes with T215V SaAS and A177S SaAS. We also report enzymological studies demonstrating that C_10_ geranyl diphosphate (GPP) can also be utilized as a substrate for the SaAS-catalyzed generation of linear and cyclic monoterpenes and monoterpene alcohols (Figure 1B). Comparisons with other sesquiterpene cyclases reveal intriguing structural differences that appear to direct cyclization reactions toward the formation of alternative major products.

## Methods

### Reagents

Unless otherwise specified, chemicals and reagents used for protein expression and purification were purchased from Fisher Scientific, Millipore Sigma, or GoldBio and used without further purification. Reagents used for crystallization were purchases from Hampton Research.

### Protein Preparation and Purification

#### Wild-type SaAS and mutants

The preparation and purification of SaAS was originally reported by Chou and colleagues,^23^ but this approach was modified to generate samples suitable for crystallization trials. Briefly, a construct of the SaAS gene in a maltose-binding protein (MBP) vector was ordered from Genscript (pET-His_6_-MBP-TEV-SaAS). A single colony was picked from BL21 (DE3) *E. coli* transformants and utilized to inoculate 6 × 1-L of 2XYT media supplemented with 50 μg/mL kanamycin. Cells were grown at 37°C with shaking at 200 RPM until the OD reached 0.5–1.0, at which point IPTG was added to a final concentration of 500 μM. Cultures were incubated at 18°C overnight, after which they were harvested by centrifugation at 6000 RPM for 20 min. Cell pellets were collected and stored at −80°C until purification.

For SaAS purification, the frozen cell pellet was resuspended in 100 mL of Buffer A [50 mM Na_2_HPO_4_ (pH 8.0), 250 mM NaCl, 10% glycerol, 1 mM tris(2-carboxyethyl)phosphine (TCEP)], also containing approximately 17 mg phenylmethylsulfonyl fluoride (PMSF, first dissolved in 1 mL EtOH), 2 pellets of EDTA-free cOmplete Mini Protease Inhibit Tablets (Roche Diagnostics), 1 mg/mL lysozyme, and 0.1 mg DNase I – grade II (Roche Diagnostics). The crude lysate mixture was sonicated using a Q700 sonicator (Qsonica) with an amplitude setting of 30% for 10 min total (cycling between 1-s on and 2-s off). The mixture was clarified by centrifugation at 18,000 RPM for 30 min. Supernatant was loaded onto a 5-mL HisTrap Ni column (Cytiva) equilibrated with Buffer A. Following a 50-mL wash step with Buffer A, the protein was eluted with a 0–100% gradient Buffer B [Buffer A plus 500 mM imidazole]. Fractions containing protein were dialyzed overnight against 1 mL of 3 mg/mL of tobacco etch virus (TEV) protease in dialysis buffer [25 mM 4-(2-hydroxyethyl)piperazine-1-ethanesulfonic acid (HEPES, pH 7.5), 250 mM NaCl, 10% glycerol, 1 mM TCEP], after which the dialyzed sample was loaded on a HisTrap Ni column with Buffer C [25 mM HEPES (pH 7.5), 250 mM NaCl, 10% glycerol, 1 mM TCEP]. Most pure protein eluted in the 30-mL wash following loading onto the column. The highest purity fractions were then injected directly onto a HiLoad 26/60 Superdex 200-pg size exclusion column (Cytiva). Fractions were analyzed by SDS-PAGE and the purest fractions were pooled and concentrated with a 10 kDa-cutoff Amicon Ultra-15 Centrifugal Filter Unit to a final concentration of approximately 8.5 mg/mL.

For the preparation of SaAS mutants, genes encoding T215V SaAS and A177F SaAS mutants were purchased directly from Genscript. The A177S, A177M, and A177I mutants were prepared utilizing the New England Biolabs (NEB) Q5^®^ site-directed mutagenesis kit with the pET-His_6_-MBP-TEV-SaAS construct. Primers for mutagenesis (Table 1) were designed by use of the NEBaseChanger tool (NEB). Mutagenesis reactions were performed as outlined by the Q5 mutagenesis kit protocol without modification. Mutant plasmids were transformed into NEB 5-α competent *E. coli* cells and were miniprepped using a QIAprep spin miniprep kit (Qiagen). Incorporation of mutations was confirmed by DNA sequencing at the University of Pennsylvania Sequencing Core and plasmids were subsequently transformed into BL21 *E. coli* cells for expression.

**Table 1.**
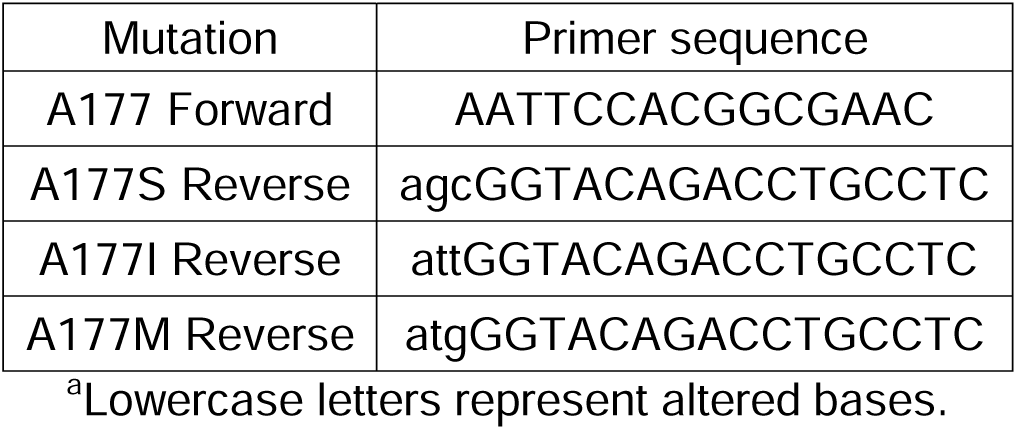
Primers for Q5 Mutagenesis^a^.

### Crystallization

SaAS crystals were obtained by the sitting drop vapor diffusion method. In a typical experiment, a 300-nL drop of protein solution [8–9 mg/mL SaAS (wild-type or mutant), 25 mM HEPES (pH 7.5), 150 mM NaCl, 10% glycerol, 1 mM TCEP] was added to 300 nL of precipitant solution [0.1 M bis-tris propane (pH 8.5–9.4), 30% PEG 6000, and 0.3–0.5% n-dodecyl-β-D-maltoside] and equilibrated against a 100-μL reservoir of precipitant solution. Diamond shaped crystals appeared in 1–2 days. Protein solution was prepared with addition of 2 mM MgCl_2_, 2 mM sodium pyrophosphate tetrahydrate, and 2 mM inhibitor. Benzyltriethylammonium chloride (BTAC) and *N,N,N*-trimethyl-1-adamantylammonium hydroxide (TMAA, TCI Chemicals) were prepared at 50 mM in HyPure^TM^ molecular biology grade water (Cytiva). *N,N*-dimethyl-1,2,3,4-tetrahydronaphthalen-2-amine hydrochloride (DMTN, Enamine) was prepared at 50 mM directly in dimethyl sulfoxide (DMSO) and *trans*-decahydroquinoline was prepared initially at 500 mM in DMSO and was diluted to 50 mM with HyPure^TM^ water. Crystals of wild-type SaAS and A177S SaAS complexed with Mg^2+^_3_-PP_i_-BTAC were grown at 21°C and all other crystals were grown at 4°C.

### Data collection and crystal structure determination

X-ray diffraction data from SaAS crystals were collected remotely at the National Synchrotron Light Source II at Brookhaven National Laboratory (Upton, NY). Data were indexed, integrated, and scaled using XDS,^25,26^ and underwent anisotropy analysis with STARANISO in the autoPROC pipeline.^27,28^ Crystals of the wild-type SaAS-Mg^2+^_3_-PP_i_-BTAC complex diffracted to 1.94 Å resolution and the initial electron density map was phased using the Phaser module of PHENIX using an AlphaFold3 model as a search model.^29,30^ Structure refinement was performed using PHENIX.refine and manual model building was performed using COOT.^31^ The refined atomic model of the wild-type SaAS-Mg^2+^_3_-PP_i_-BTAC complex was used as search model to phase initial electron density maps for all remaining structures described herein.

After initial refinement of each structure, residue side chains with limited or no electron density in 2m|*F_o_*|-D|*Fc*| maps were truncated or removed. Solvent molecules were placed manually in peaks >3σ in the |*F_o_*|-|*F_c_*| map. During the final stage of refinement, ligands and other non-proteogenic molecules were added to the model. Validation of all final models was achieved with Molprobity.^32^ Data collection and refinement statistics are listed in Table 1. Figures of structures were prepared with PyMOL (PyMOL Molecular Graphics System, version 2.1, Schrödinger, LLC). Active site contours were generated using the GetCleft component of the NRGsuite plug-in for PyMOL.^33^

### Enzyme Activity and Product Analysis

Gas chromatography-mass spectrometry (GC-MS) was utilized for identification and quantification of reaction products with an Agilent 8890 GC/5597C MSD system equipped with a J&W HP-5ms Ultra Inert GC capillary column (30 m x 0.25 mm x 0.25 µm). Geranyl diphosphate (GPP), farnesyl diphosphate (FPP), and geranylgeranyl diphosphate (GGPP) for all activity and product profiling experiments were purchased from Isoprenoids.com and resuspended in 7:3 methanol:10 mM NH_4_HCO_3_ at a concentration of 10 mM for GPP and FPP and 5 mM for GGPP. The temperature program for all samples was initiated with an oven temperature of 60°C held for 2 min followed by a ramp to 240°C at 20°C/min. Following a solvent delay of 3 min, positive EI mode was used to collect MS data. Some products were identified and confirmed by comparison of retention times and mass spectra with those of authentic commercial standards. Mass spectra were also utilized for identification by comparison to spectra archived in the National Institute of Standards and Technology (NIST) database for further confirmation.

#### Relative Activity

Activity measurements were performed in triplicate on a 200-µL scale using an enzyme concentration of 200 nM in reaction buffer [25 mM HEPES (pH 7.5), 150 mM NaCl, 1 mM TCEP, 10% glycerol] and 1.5 mM MCl_2_ (M = Co^2+^, Mg^2+^, Mn^2+^, Ni^2+^). Reactions were initiated by addition of 10 µL of a 200 µM substrate stock and left at room temperature for four different time points. Ethyl acetate containing a hexadecane internal standard was utilized to quench reactions with vortexing for 3 × 10-s cycles. 100 µL of the ethyl acetate phase was removed and transferred to a separate vial for GC-MS analysis. To measure enzyme activity, samples were run with an injection volume of 4 µL using selected ion monitoring (SIM) for ions at the following m/z: 57.10, 71.00, 77.00, 79.10, 85.10, 91.00, 93.00, 121.00, 136.20, 226.20. Peaks at 9.74 min were integrated and normalized by comparison to the internal standard peak at 9.19 min.

#### Product profiles

Enzyme products were identified and quantified using a 200-µL scale with 5 µM enzyme and 400 µM substrate in reaction buffer [25 mM HEPES (pH 7.5), 150 mM NaCl, 1 mM TCEP, 10% glycerol] and 1.5 mM MCl_2_ (M = Co^2+^, Mg^2+^, Mn^2+^, Ni^2+^). Reactions were initiated by addition of substrate immediately followed by layering of 200 µL of hexanes over the aqueous phase. Generally, reactions were incubated overnight at room temperature after which the reaction mixture was vortexed for 3 × 10-s cycles and then transferred to a 1.5 mL Eppendorf tube which was then spun down at 12,000 rpm for 15 seconds. 100 µL of the organic phase would then be transferred to a separate GC vial for analysis with GC-MS. Samples for product profile determination were run with a 1-µL injection volume using GC-MS as described above.

## Results

### Enzyme Activity and Product Arrays

Upon its discovery, SaAS was reported to be a moderate-fidelity sesquiterpene cyclase, generating avermitilol as 85% of total products detected in the presence of Mg^2+^.^23^ Our measurements indicate that SaAS is in fact a high-fidelity cyclase in the presence of Mg^2+^, generating avermitilol as 95.5% of total products detected (Table 2). Significant variations in product profiles result when different metal ion cofactors are utilized (Figure 2). Relative activity is highest with Mn^2+^, although Co^2+^ and Mg^2+^ still confer high levels of activity only reduced by approximately 2.4- and 1.6-fold respectively. Activity is reduced by approximately 16-fold with Ni^2+^. Class I terpene cyclases are often quite selective for a particular divalent metal ion (e.g., usually Mg^2+^, as observed for bacterial cyclases,^34^ but sometimes Mn^2+^), so it is interesting that SaAS exhibits high levels of activity with three different metal ions – Mg^2+^, Mn^2+^, and Co^2+^.

**Table 2.**
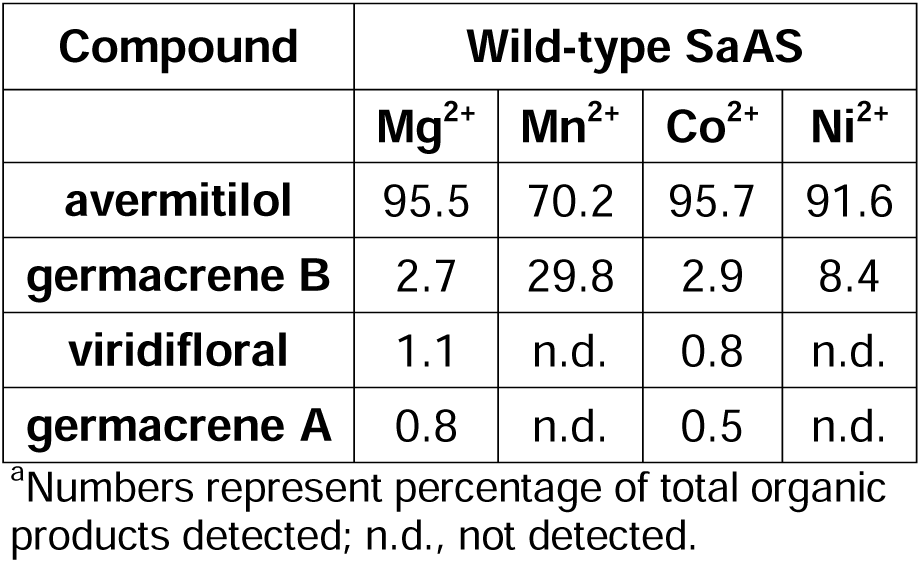
FPP cyclization products generated with different metal ions^a^.

**Figure 2.**
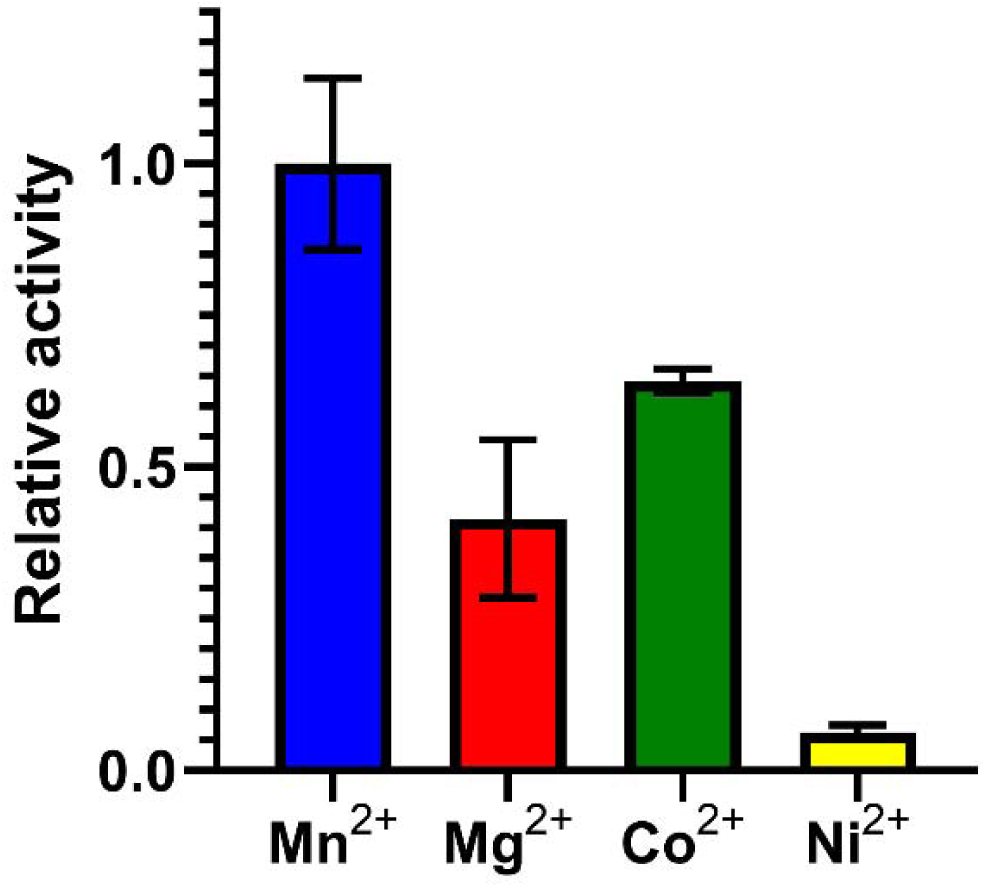
Influence of divalent metal ions on SaAS activity. Error bars indicate standard deviation for 3 technical replicates.

**Table 2.** Data Collection and Refinement Statistics.

| Structure | SaAS Mg<sup>2+</sup><sub>3</sub>-PP<sub>i</sub>- | T215V SaAS Mg<sup>2+</sup><sub>3</sub>- | A177S SaAS Mg<sup>2+</sup><sub>3</sub>- | SaAS Mg<sup>2+</sup><sub>3</sub>-PP<sub>i</sub>- | SaAS Mg<sup>2+</sup><sub>3</sub>- |
| --- | --- | --- | --- | --- | --- |

**Table 2. Data Collection and Refinement Statistics**
|  | BTAC | PP<sub>i</sub>-BTAC | PP<sub>i</sub>-BTAC | TMAA | PP<sub>i</sub>-DMTN |
| --- | --- | --- | --- | --- | --- |
| **Unit Cell** |  |  |  |  |  |
| space group | C2 | C2 | C2 | C2 | C2 |
| *a*, *b*, *c* (Å) | 95.53, 56.33,<br>72.64 | 95.37, 56.50,<br>72.56 | 95.571, 56.10,<br>72.69 | 95.552, 56.25,<br>72.77 | 95.55, 56.37,<br>76.12 |
| $\alpha$, $\beta$, $\gamma$ (deg) | 90, 125.15, 90 | 90, 125.15, 90 | 90, 125.34, 90 | 90, 125.3, 90 | 90, 125.31, 90 |
| **Data Collection** |  |  |  |  |  |
| resolution (Å) | 1.28 | 1.42 | 1.30 | 1.42 | 1.44 |
| total/unique no. of reflections | 213,848/67,371 | 198,989/59,408 | 322,032/73,659 | 270,897/57,615 | 217,904/57,031 |
| $R_{\text{merge}}^{a,b}$ | 0.084 (0.634) | 0.083 (1.053) | 0.076 (0.912) | 0.104 (1.142) | 0.110 (1.198) |
| $R_{\text{pim}}^{a,c}$ | 0.053 (0.537) | 0.052 (0.736) | 0.039 (0.652) | 0.051 (0.628) | 0.063 (0.746) |
| $CC_{1/2}^{a,d}$ | 0.992 (0.435) | 0.998 (0.344) | 0.998 (0.332) | 0.997 (0.457) | 0.997 (0.320) |
| $I/\sigma(I)^a$ | 8.1 (1.5) | 8.5 (1.1) | 9.2 (0.9) | 7.9 (1.2) | 7.7 (1.2) |
| redundancy<sup>a</sup> | 3.2 (1.8) | 3.3 (2.8) | 4.4 (2.4) | 4.7 (3.9) | 3.8 (3.4) |
| completeness (%)<sup>a</sup> | 82.3 (38.5) | 99.5 (97.4) | 94.4 (59.0) | 96.4 (99.9) | 99.3 (95.3) |
| **Refinement** |  |  |  |  |  |
| reflections used in refinement/test set | 67,071/1,991 | 58,775/2,962 | 73,587/2,000 | 57,296/1,990 | 56,103/2,766 |
| $R_{\text{work}}^{a,e}$ | 0.196 (0.302) | 0.184 (0.298) | 0.179 (0.329) | 0.160 (0.290) | 0.175 (0.259) |
| $R_{\text{free}}^{a,e}$ | 0.213 (0.313) | 0.202 (0.309) | 0.195 (0.347) | 0.200 (0.336) | 0.197 (0.286) |
| no. of protein chains | 1 | 1 | 1 | 1 | 1 |
| no. of nonhydrogen atoms | 2,737 | 2,766 | 2,707 | 2,750 | 2,788 |
| protein | 2,529 | 2,525 | 2,495 | 2,530 | 2,536 |
| ligand | 30 | 34 | 16 | 34 | 20 |
| solvent | 178 | 207 | 196 | 186 | 232 |
| average *B* factor (Å<sup>2</sup>) | 17 | 17 | 16 | 20 | 15 |
| protein | 17 | 17 | 15 | 19 | 15 |
| ligand | 16 | 18 | 13 | 22 | 14 |
| solvent | 22 | 23 | 23 | 26 | 21 |
| root-mean-square deviation from ideal geometry |  |  |  |  |  |
| bonds (Å) | 0.005 | 0.005 | 0.005 | 0.005 | 0.005 |
| angles (deg) | 0.8 | 0.8 | 0.8 | 0.8 | 0.8 |
| Ramachandran plot<sup>f</sup> |  |  |  |  |  |
| Favored (%) | 99.09 | 99.08 | 99.69 | 99.10 | 99.39 |
| Allowed (%) | 0.91 | 0.92 | 0.31 | 0.90 | 0.61 |
| Outliers (%) | 0.00 | 0.00 | 0.00 | 0.00 | 0.00 |
| Molprobit score<sup>f</sup> | 0.80 | 0.90 | 0.65 | 0.80 | 0.64 |
| PDB entry | 9Y6I | 9Y6K | 9Y6L | 9Y6M | 9Y6J |

<sup>a</sup>Values in parentheses refer to the highest-resolution shell of data.
<sup>b</sup> $R_{\text{merge}} = \sum_h \sum_i |I_{h,i} - \langle I \rangle_h| / \sum_h \sum_i I_{h,i}$ , where $\langle I \rangle_h$ is the average intensity calculated for reflection *h* from *i* replicate measurements.
<sup>c</sup> $R_{\text{p.i.m.}} = (\sum_h (1/(N-1))^{1/2} \sum_i |I_{h,i} - \langle I \rangle_h|) / \sum_h \sum_i I_{h,i}$ , where *N* is the number of reflections and $\langle I \rangle_h$ is the average intensity calculated for reflection *h* from replicate measurements.
<sup>d</sup>Pearson correlation coefficient between random half-datasets.
<sup>e</sup> $R_{\text{work}} = \sum ||F_o| - |F_c|| / \sum |F_o|$ for reflections contained in the working set. $|F_o|$ and $|F_c|$ are the observed and calculated structure factor amplitudes, respectively. $R_{\text{free}}$ is calculated using the same expression for reflections contained in the test set held aside during refinement.
<sup>f</sup>Calculated with MolProbity
Swapped the order to match text.

Metal-dependent product profile arrays were also investigated. Only subtle changes are observed when comparing the product arrays generated using Co^2+^ or Ni^2+^ compared with Mg^2+^ (Table 2). However, the array is notably altered when Mn^2+^ is substituted for Mg^2+^, such that the percentage of avermitilol generated is reduced by 25% accompanied by an 11-fold increase in the generation of germacrene B (Table 2). Premature derailment of the germacryl cation in the normal cyclization cascade leads to germacrene B. Although enzyme activity is highest with Mn^2+^, it is notable that increased activity is accompanied by increased cyclization promiscuity.

The substrate scope of SaAS was briefly investigated by testing GPP and GGPP for activity. When incubated with GGPP, SaAS does not generate any diterpene products. Presumably, the larger C_20_ substrate GGPP does not readily fit in an active site pocket that normally accommodates the smaller C_15_ substrate FPP. However, when incubated with GPP SaAS generates an array of monoterpene products, including linear and cyclic hydrocarbons along with alcohol and ether products (Table 3). Notably, reactions leading to products such as limonene and 1,8-cineole require a diphosphate ionization-recombination-reionization sequence to enables formation of a 6-membered ring product. This chemistry is unrelated to that of the native FPP cyclization reaction catalyzed by SaAS, which does not proceed through a diphosphate ionization-recombination-reionization sequence.

**Table 3.**
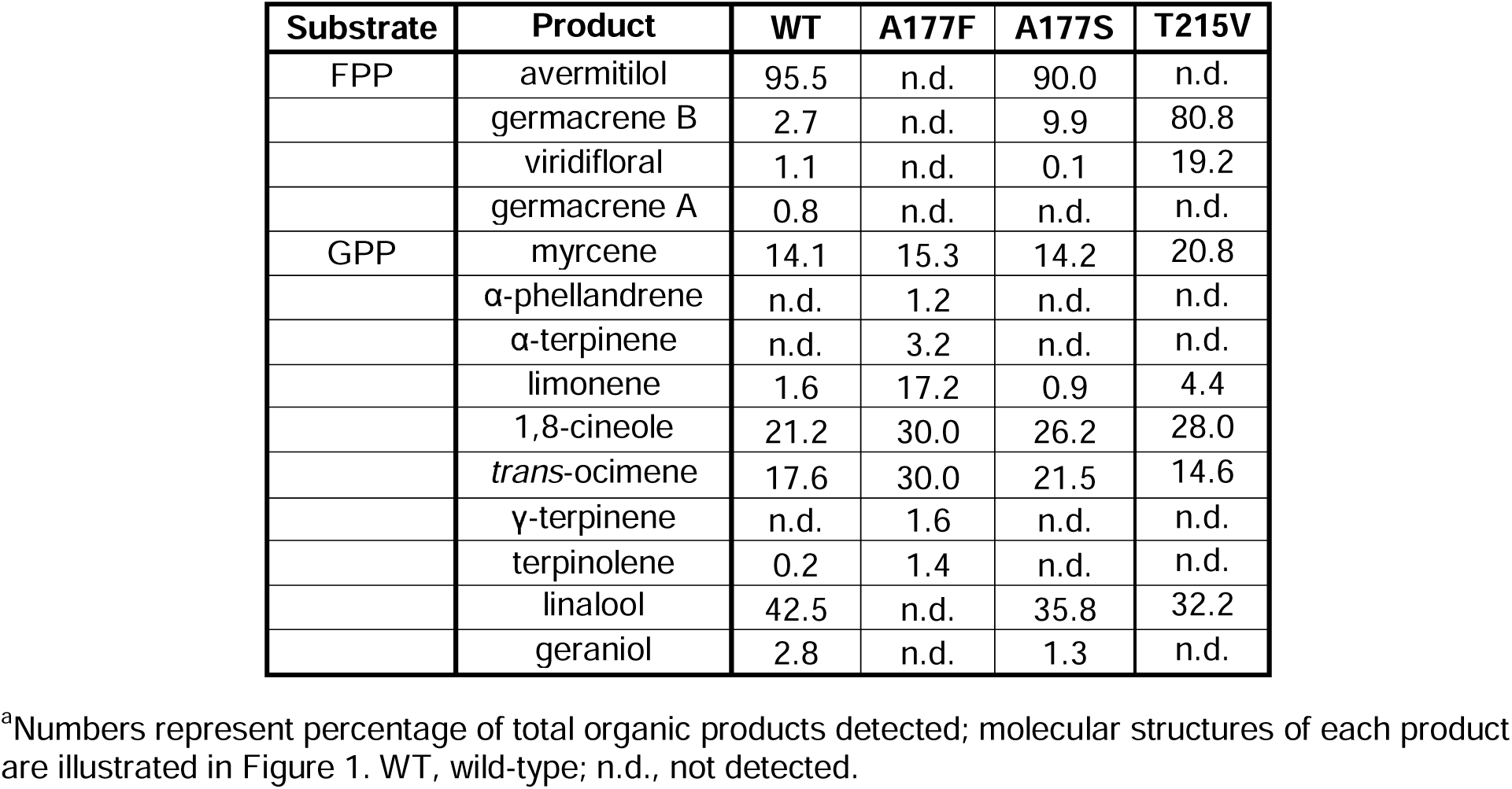
Sesquiterpene and Monoterpene Product Arrays^a^.

As a first step in probing the functional role of residues lining the active site pocket, site-directed mutagenesis was employed to substitute larger side chains for A177. While A177S is catalytically active and generates a product array similar to that of the wild-type enzyme (Table 3), the A177I and A177M mutants exhibit no activity with FPP or GPP. Thus, significantly larger amino acids substituted for A177 will hinder productive binding of the substrate. Interestingly, however, A177S SaAS and A177F SaAS can utilize GPP as a substrate to generate somewhat altered monoterpene product arrays (Table 3). Monoterpene alcohol products are not detected, but the mechanism leading to the monoterpene ether 1,8-cineole proceeds through an alcohol intermediate.

### Structure of the SaAS-Mg^2+^_3_-PP_i_-BTAC Complex

SaAS shares the canonical class I terpene cyclase fold with helices A–K associated so as to form an active site pocket in the center of a helical bundle. The crystal structure of the SaAS-Mg^2+^_3_-PP_i_-BTAC complex reveals a fully closed active site conformation, with the Hα-1 loop and the J-K loop capping the active the site (Figure 3A). The transition from open to closed active site conformations in a class I terpene cyclase is driven by Mg^2+^_3_-PP_i_ complexation and serves to protect carbocation intermediates in catalysis from premature quenching by bulk solvent.^2^ SaAS crystallizes as a dimer (Figure 3B), but the functional relevance of quaternary structure is unclear since cooperativity is not observed in steady-state kinetics.^23^ Even so, the PISA server^35^ predicts a biological dimer with 1702 Å^2^ total buried surface area.

**Figure 3.**
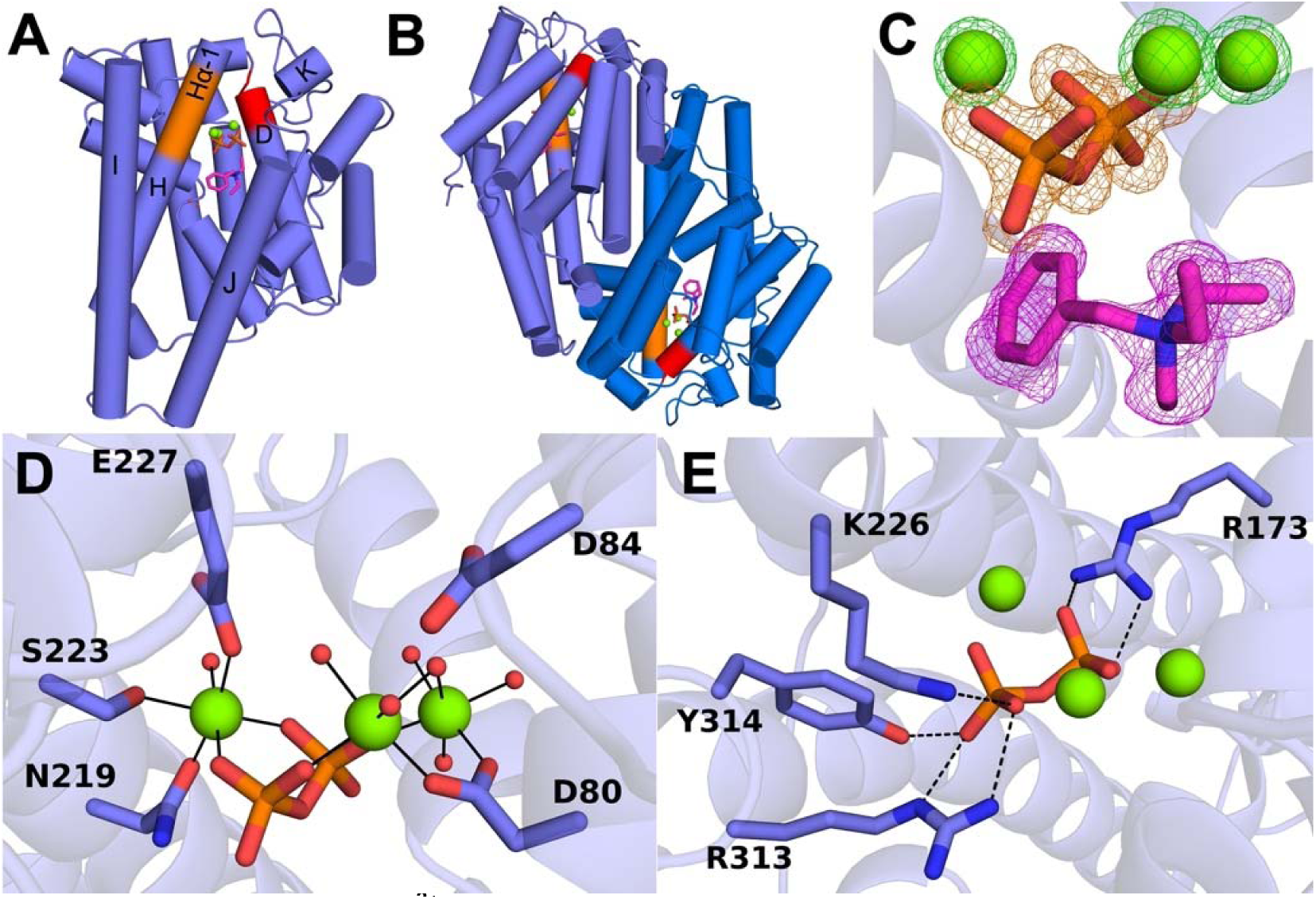
Structure of the SaAS-Mg^2+^_3_-PP_i_-BTAC complex. (A) SaAS adopts the conserved α fold of a class I terpene cyclase. The bundle of helices A–K surround the active site pocket, where 3 Mg^2+^ ions (green spheres), the PP_i_ anion (stick figure, P = orange, O = red), and BTAC (stick figure, C = magenta, N = blue) are bound. Aspartate-rich and NSE/DTE metal binding motifs are red and orange, respectively. (B) The dimer interface of SaAS corresponds to that commonly observed in other dimeric class I sesquiterpene cyclases and buries a total of 1702 Å^2^ surface area. (C) Polder omit maps of the Mg^2+^ ions, PP_i_, and BTAC contoured at 5.5σ, 6σ, and 4σ, respectively. (D) Mg^2+^ coordination polyhedra; metal coordination interactions are represented by thin solid lines, and water molecules are represented as small red spheres. (D) The PP_i_ anion accepts hydrogen bonds (dashed black lines) from three basic residues and a tyrosine.

The electron density map reveals clear density corresponding to the Mg^2+^_3_-PP_i_ cluster and a single molecule of BTAC in the active site pocket (Figure 3C). Metal binding motifs conserved in class I terpene synthases are found as the aspartate-rich motif D^80^DXXD^84^ and the NSE/DTE motif N^219^S^223^E^227^, each of which is involved in binding the Mg^2+^_3_ cluster (Figure 3D). In addition to metal coordination interactions, the PP_i_ anion forms hydrogen bonds with R313, Y314, K226, and R173 (Figure 3E).

BTAC binding is accommodated by several interactions in the active site pocket. The aromatic ring centroid of F77 is 4.4 Å away from the quaternary ammonium nitrogen of BTAC, indicative of a strong cation-π interaction (Figure 4A). The phenyl ring of BTAC additionally makes van der Waals contacts with aromatic residues Y304 and W300. Comparisons of the SaAS-Mg^2+^_3_-PP_i_-BTAC complex with other sesquiterpene cyclase-Mg^2+^_3_-PP_i_-BTAC complexes show that each enzyme positions and orients the Mg^2+^_3_-PP_i_ complex similarly, but the orientation of BTAC varies from one enzyme to another, seemingly rotating around the quaternary ammonium nitrogen (Figure 4B). Variations in BTAC binding reflect variations in active site contours in different sesquiterpene cyclases.

**Figure 4.**
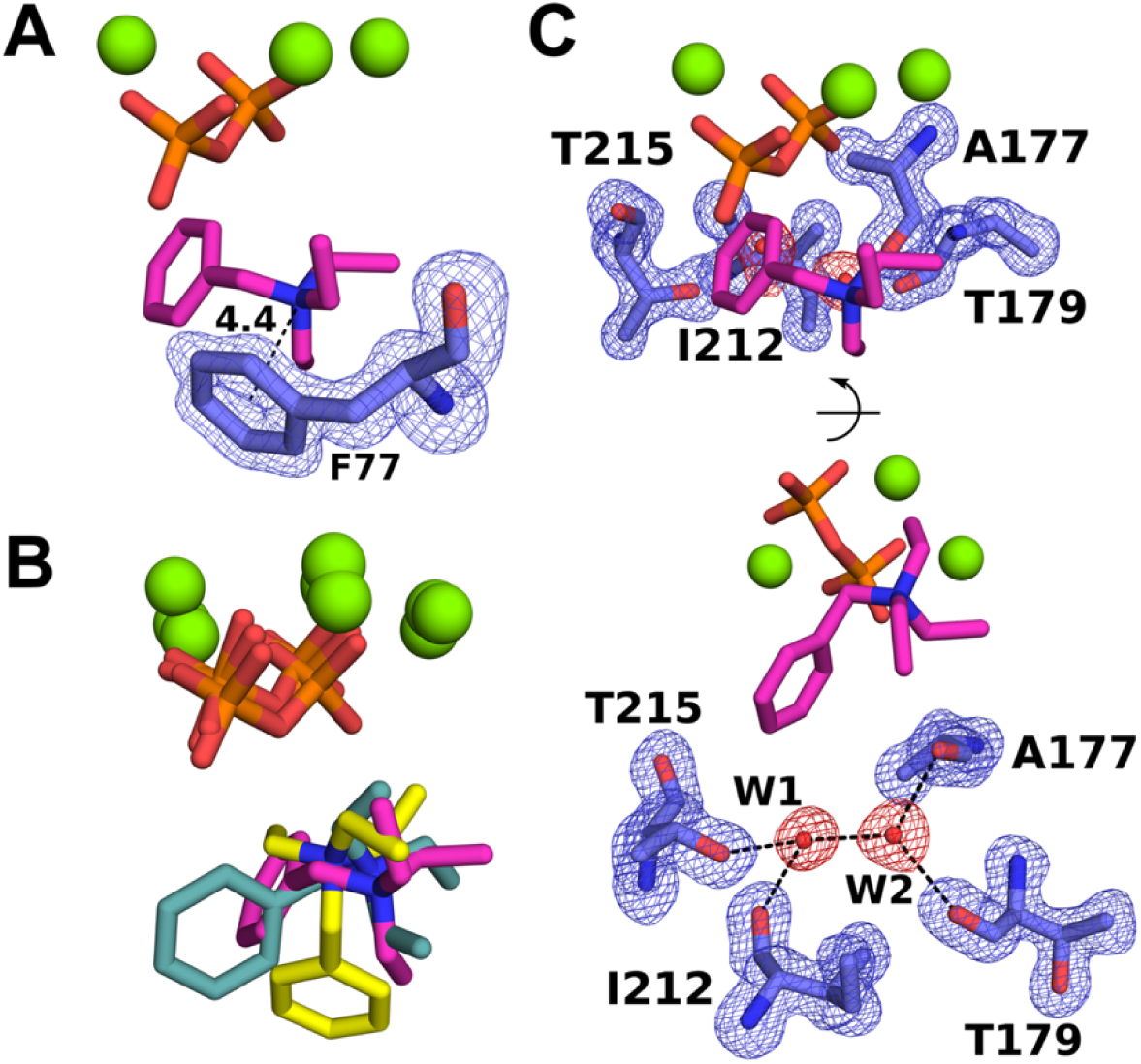
Interactions of BTAC in the SaAS-Mg^2+^_3_-PP_i_-BTAC complex. (A) Polder omit map of F77 showing a cation-π interaction between F77 and the quaternary ammonium group of BTAC. (B) Alignment of sesquiterpene cyclase-BTAC complexes reveals similar positioning of the Mg^2+^_3_-PP_i_ cluster, but alternate orientations for BTAC related by rotations around the quaternary ammonium nitrogen atom. Enzyme residues are not shown for clarity; BTAC positions correspond to SaAS (magenta), *epi*-isozizaene synthase (EIZS, teal), and presilphiperfolan-8β-ol synthase (BcBOT2, yellow). (C) Polder omit maps of protein residues and bound water molecules in the active site contoured at 4σ. (C) Hydrogen bond network involving active site residues and two trapped water molecules, W1 and W2.

Also notable is the binding of two hydrogen-bonded water molecules in a distinctly polar crevice in the active site pocket, held in place by hydrogen bonds with enzyme residues (Figure 4C). The side chain of T215 and the backbone carbonyl of I212 form hydrogen bonds with water molecule W1, and the backbone carbonyl groups of A177 and T179 form hydrogen bonds with water molecule W2. It is not often that water molecules are observed in the enclosed active site pocket of a terpene cyclase in the closed conformation. However, since water is a co-substrate in the reaction generating avermitilol, the active site has evolved to manage and manipulate water as well as the isoprenoid substrate.

### Structure of the A177S SaAS-Mg^2+^_3_-PP_i_-BTAC Complex

The residue corresponding to A177 in SaAS has been subject to mutagenesis in other sesquiterpene cyclases to probe its influence on hydroxylation chemistry.^36^ This prompted our interest in the study of site-specific mutations at this position. The product profile of A177S SaAS is relatively unchanged from that of the wild-type enzyme (Table 3). The crystal structure of the A177S SaAS-Mg^2+^_3_-PP_i_-BTAC complex was determined at 1.30 Å resolution, revealing that active site water molecules W1 and W2 remain in place (Figure 5A,C); the newly introduced hydroxyl group of S177 forms a weakly polar interaction with water molecule W2. In wild-type SaAS the backbone carbonyl groups of A177 and T179 form hydrogen bonds with water W2, and these hydrogen bonds are conserved in A177S SaAS (Figure 5B). Comparable enzyme activities measured for A177S SaAS and wild-type SaAS (Figure 2) are consistent with a conserved position for a catalytically-required water molecule in these enzymes.

**Figure 5.**
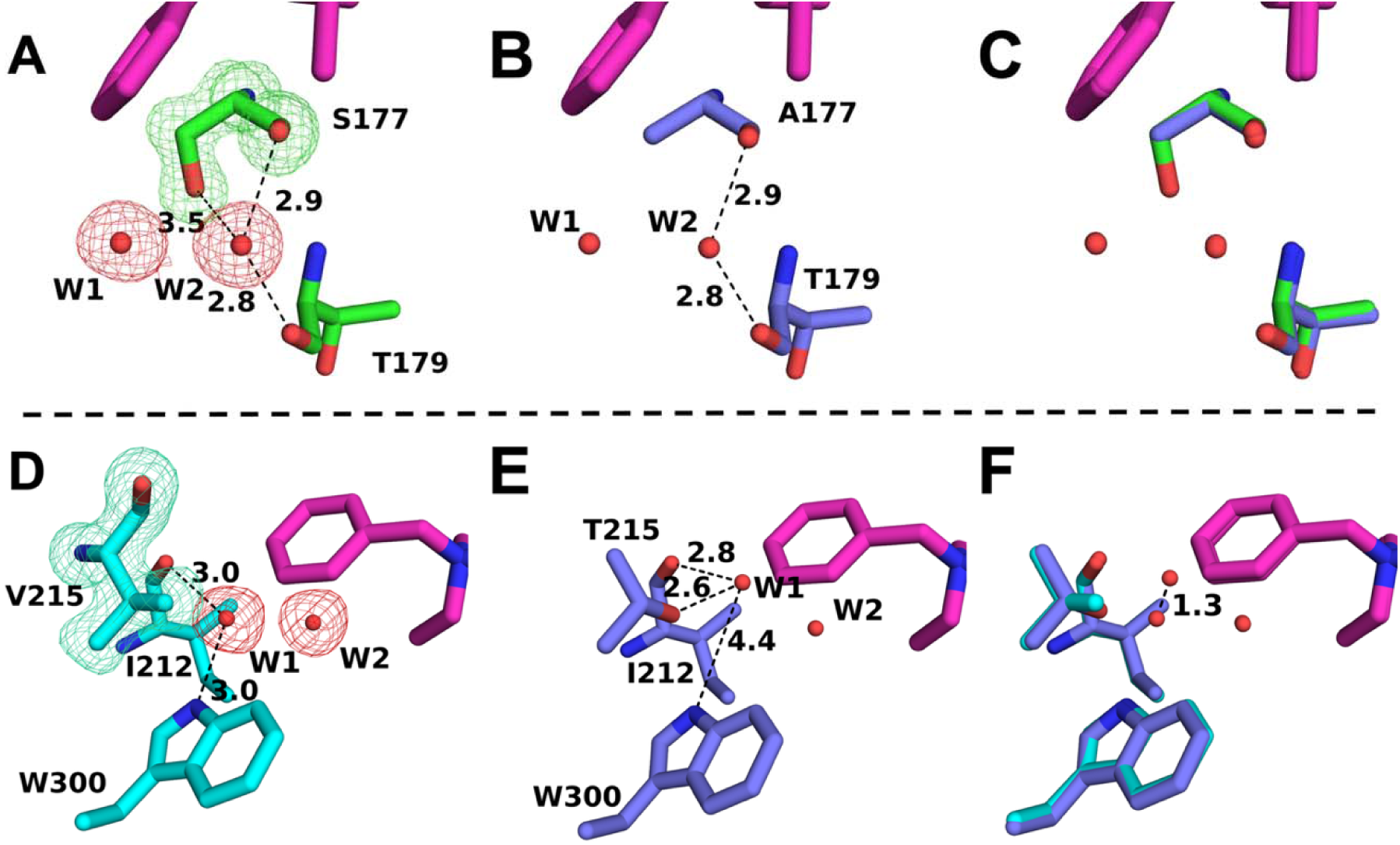
Crystal structures of A177S SaAS and T215V SaAS. (A) Polder omit map of the mutated residue in A177S SaAS and active site water molecules W1 and W2 (contoured at 4σ) reveals a 3.5 Å-long weakly polar interaction between the S177 hydroxyl group and water molecule W1. (B) Hydrogen bond interactions with water W2 from the backbone carbonyl groups of A177 and T179 in wild-type SaAS; these interactions are conserved in A177S SaAS. (C) Overlay of wild-type SaAS and A177S SaAS showing essentially identical positioning of water molecules W1 and W2. (D) Polder omit map of V215 (contoured at 4σ) in the polar active site crevice of T215V SaAS. Hydrogen bond interactions with water molecule W1 are indicated by dashed black lines. (E) Polar active site crevice of wild-type SaAS; interactions with water molecule W1 are indicated by dashed black lines. Comparison with the wild-type enzyme (E) shows that water molecule W1 shifts 1.3 Å from its position in the wild-type enzyme to form a hydrogen bond with W300, as highlighted in the overlay of wild-type SaAS and T215V SaAS (F).

### Structure of the T215V SaAS-Mg^2+^_3_-PP_i_-BTAC Complex

To further probe the functional importance of T215 as it forms a hydrogen bond with water W1, we hypothesized that mutation of T215 might impact product outcome if water management were compromised by the mutation. Indeed, T215V SaAS generates no avermitilol when incubated with FPP, but instead generates viridifloral along with greater quantities of germacrene B, a premature elimination product (Table 3). The X-ray crystal structure of the T215V SaAS-Mg^2+^_3_-PP_i_-BTAC complex reveals that water molecule W1 moves 1.3 Å from its position in the wild-type enzyme complex to form a hydrogen bond with W300 (Figure 5D,E). Overlay of the wild-type and T215V structures shows that W1 moves 1.3 Å while the position of W2 remains constant (Figure 5F). The indole nitrogen of W300 forms a hydrogen bond interaction with W1 in place of T215. The 1.3 Å-shift of W1 completely disrupts the ability to generate avermitilol, suggesting that W1 is the catalytically-required water molecule that reacts with bicyclogermacrene to generate avermitilol in the final step of catalysis, as shown in Figure 1.

### Structure of the SaAS-Mg^2+^_3_-PP_i_-TMAA Complex

To further study the binding of cationic ligands capable of mimicking carbocation binding and stabilization, we cocrystallized SaAS with *N,N,N*-trimethyl-1-adamantyl-ammonium hydroxide (TMAA, Figure 1) and determined its structure at 1.42 Å resolution. The adamantane skeleton is accommodated well in the enzyme active site and electron density for the entire ligand is well resolved (Figure 6A). The positively charged quaternary ammonium group forms a cation-π interaction with F77 (Figure 6B), as also observed for BTAC (Figure 4A). The binding mode of TMAA is similar to that of BTAC with only a slight shift in the positioning of the ammonium group and same relative positioning of the adamantane skeleton compared with the benzyl group of BTAC (Figure 6C).

**Figure 6.**
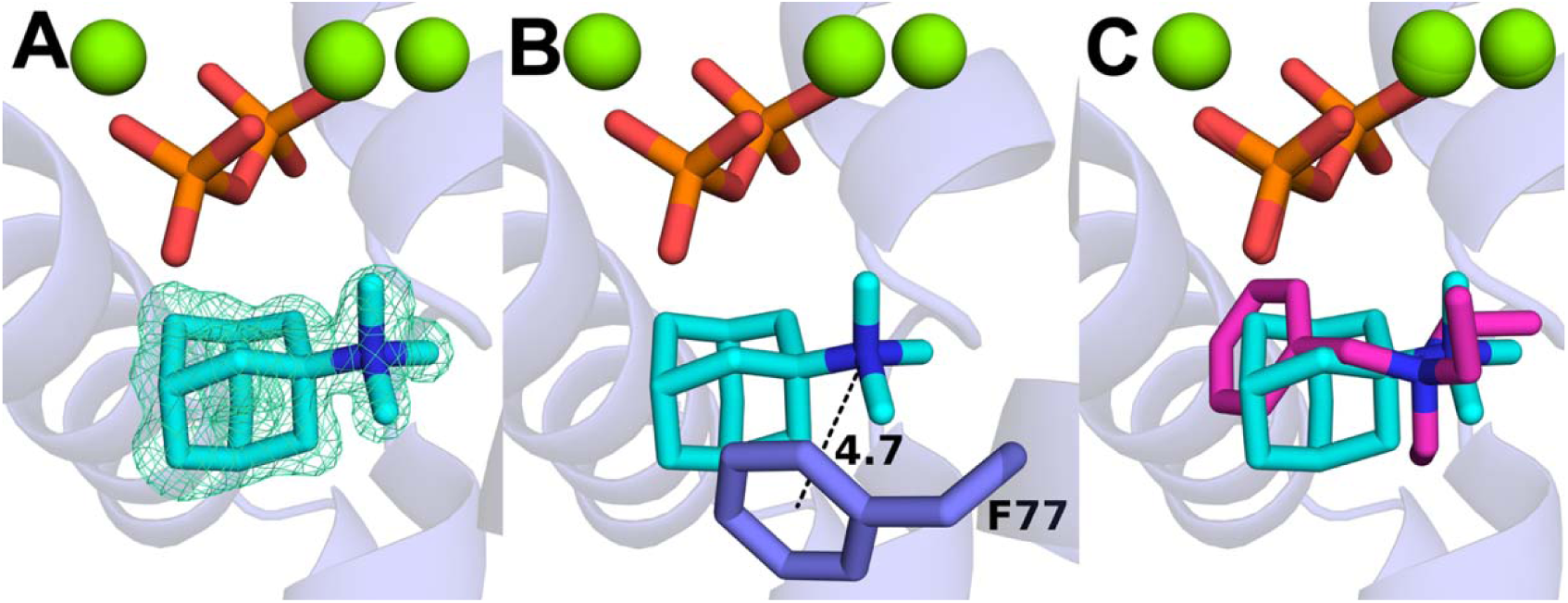
Structure of the SaAS-Mg^2+^_3_-PP_i_-TMAA Complex. (A) Polder omit map of TMAA contoured at 6σ. (B) Cation-π interaction formed between F77 and the positively charged quaternary ammonium group of TMAA. (C) Overlay of the SaAS-TMAA complex and the SaAS-BTAC complex showing similar positions for the positively charged ammonium group. Water molecules are omitted for clarity.

### Structure of the SaAS-Mg^2+^_3_-PP_i_-DMTN Complex

To study the binding of a protonated tertiary ammonium group similarly capable of mimicking aspects of carbocation binding and stabilization, we cocrystallized SaAS with a bicyclic tetralin derivative, *N,N*-dimethyl-1,2,3,4-tetrahydronaphthalen-2-amine hydrochloride (DMTN, Figure 1), and determined the structure of the complex at 1.44 Å resolution. The bicyclic amine binds deep in the active site pocket and its presumably-protonated tertiary ammonium group donates a hydrogen bond to the PP_i_ anion (Figure 7A,B); the ammonium nitrogen atom of DMTN is positioned similarly to that of BTAC (Figure 7C). Comparisons of the SaAS complexes with BTAC, TMAA, and DMTN reveals that the binding orientation of diverse ammonium ion derivatives is driven by favorable charge-charge interactions with the PP_i_ anion and cation-π interactions with the aromatic ring of F77. There is no particular difference evident in the binding of tertiary versus quaternary ammonium groups.

**Figure 7.**
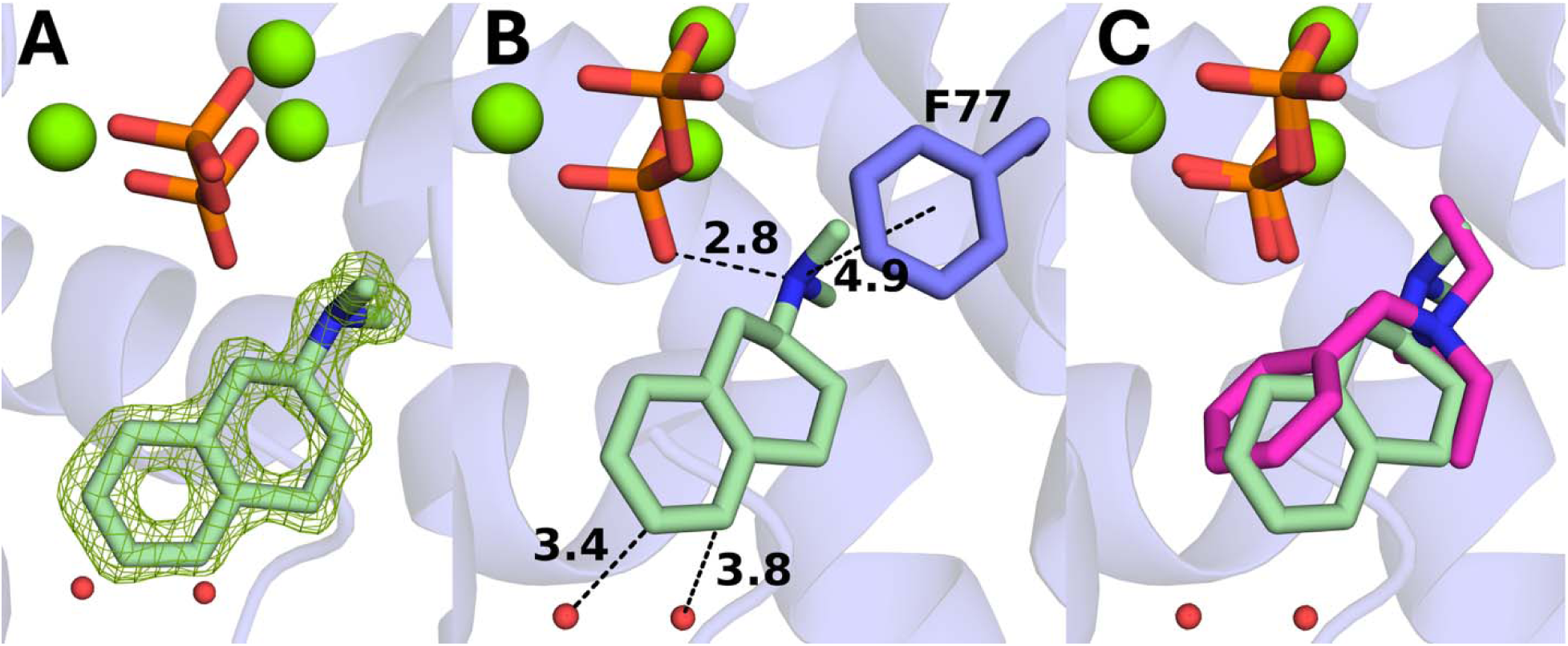
Structure of the SaAS-Mg^2+^_3_-PP_i_-DMTN Complex. (A) Polder omit map of DMTN contoured at 6σ. (B) Key interactions of DMTN with the PP_i_ anion, F77, and trapped water molecules W1 and W2. (C) Overlay of the SaAS complexes with DMTN (olive) and BTAC (magenta) reveals that the positively charged ammonium groups bind similarly regardless of whether they are quaternary or tertiary, but the tertiary ammonium group is less sterically hindered and can get close enough to the PP_i_ anion to donate a hydrogen bond.

## Discussion

Examples of terpene cyclases with the ability to utilize different divalent metal ions for catalysis are relatively rare, with bacterial sesquiterpene cyclases usually being selective for Mg^2+^.^34^ In some cases, however, class I terpene cyclases can function with alternative divalent metal ions. For example, sabinene synthase from *Thuja plicata* can function with Mn^2+^, Mg^2+^, Co^2+^, and Ni^2+^, and these metal ions yield generally similar product profiles.^37^ Intriguingly, some terpene cyclases exhibit alternative product arrays depending on the identity of the divalent metal ion. For example, the class I terpene cyclase OILTS from a giant virus generates (+)-germacrene D-4-ol with Mg^2+^ or Zn^2+^, but generates (+)-cubebol with Mn^2+^, Ni^2+^, or Co^2+^.^38^

In SaAS, the highest level of relative activity is achieved with Mn^2+^ (Figure 2), but the product profile varies significantly depending on the metal ion utilized (Table 2). Curiously, the use of Mn^2+^ instead of Mg^2+^ increases the generation of the early elimination product germacrene B. Not only the metal ion, but also the protein ligands to a metal ion can influence the product profile. For example, mutagenesis of metal-binding motifs in trichodiene synthase from *Fusarium sporotrichioides* results in changes to the product profile.^39^ Slight differences in the ionic radii of different metal ions, or slight changes in the positioning of metal ions resulting from mutagenesis of metal ligands, likely reposition the substrate diphosphate group so as to influence reaction pathways.

It is notable that SaAS exhibits robust activity using the smaller isoprenoid substrate GPP. When incubated with GPP, SaAS generates a wide array of linear, cyclic, and hydroxylated monoterpene products (Figure 1, Table 3). Sometimes, but not always, a cyclase that utilizes a larger isoprenoid substrate will also be able to utilize a smaller isoprenoid substrate. For example, the diterpene cyclase venezuelaene A from *Streptomyces venezuelae* ATCC 15439 utilizes GGPP to generate venezuelaene A, but it can also utilize FPP to generate germacrene A or hedycaryol, or GPP to generate geraniol.^40^

SaAS employs strikingly different chemistry for GPP cyclization compared with FPP cyclization (Figure 1). The generation of cyclic monoterpenes from GPP requires a specific ionization-recombination-reionization sequence to generate the core 6-membered ring; such chemistry does not occur in the SaAS-catalyzed reaction with FPP, but it does occur in sesquiterpene cyclases that generate initial 6-membered rings from FPP in their cyclization cascades, such as *epi*-isozizaene synthase.^11^ The three-dimensional contour of the active site plays an important role in enabling and directing cyclization chemistry, particularly with regard to the initial cyclization step of catalysis and the size of the substrate that can be accommodated in the active site (Figure 8). The production of 1,8-cineole by SaAS is also notable since the reaction pathway leading to this bicyclic ether proceeds through a sesquiterpene alcohol intermediate – water molecules W1 and/or W2 trapped in the active site likely serve as the source of the hydroxyl group for this intermediate, and thence the ether oxygen atom of 1,8-cineole.

**Figure 8.**
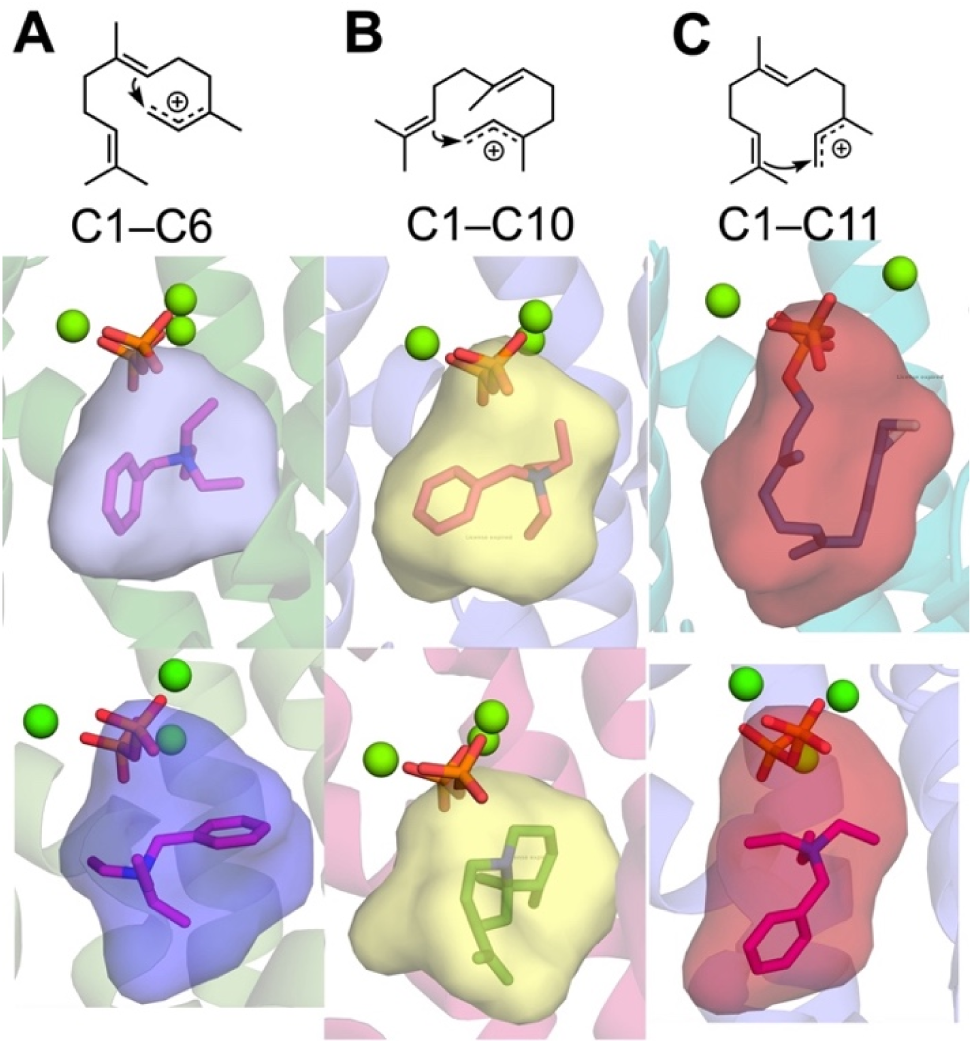
Terpene cyclase active site contours and initial cyclization steps. (A) C1-C6 cyclization step directed by *epi*-isozizaene synthase (top, PDB 3KB9) and trichodiene synthase (bottom, PDB 2Q9Y). (B) C1-C10 cyclization step directed by SaAS (top, PDB 9Y6I) and aristolochene synthase (bottom, PDB 4KVY). (C) C1-C11 cyclization (anti-Markovnikov) directed by pentalenene synthase (top, 6WKD) and presilphiperfolan-8β-ol synthase (bottom, 8H6U).

A terpene cyclase contain a distinct active site contour that guides reactive substrate conformations toward a specific product outcome. Structural studies of terpene cyclases have been crucial for understanding such mechanistic features. In particular, structures of enzymes complexed with catalytic metal ions, the PP_i_ co-product, and substrate analogues are exceedingly useful for understanding structure-function relationships. Most often, such complexes reveal conformational changes that enclose the active site, so as to protect carbocation intermediates in catalysis from premature quenching by exposure to bulk solvent. This is not to say that terpene cyclases cannot engage in biosynthetic reactions with water; to the contrary, terpene cyclases are indeed capable of managing and manipulating a trapped water molecule to quench a specific carbocation intermediate (Figure 9). The likely utilization of water molecule W1 by SaAS to generate avermitilol or side product viridifloral serves as a prominent example of a water management strategy in terpene biosynthesis. The T215V substitution blocks avermitilol formation, but this substitution enhances the generation of viridifloral (as well as germacrene B); enhanced viridifloral generation appears to result from the repositioning of W1 so that it cannot engage in anti-Markovnikov addition, but rather Markovnikov addition to the reactive C=C bond of bicyclogermacrene (Figure 9).

**Figure 9.**
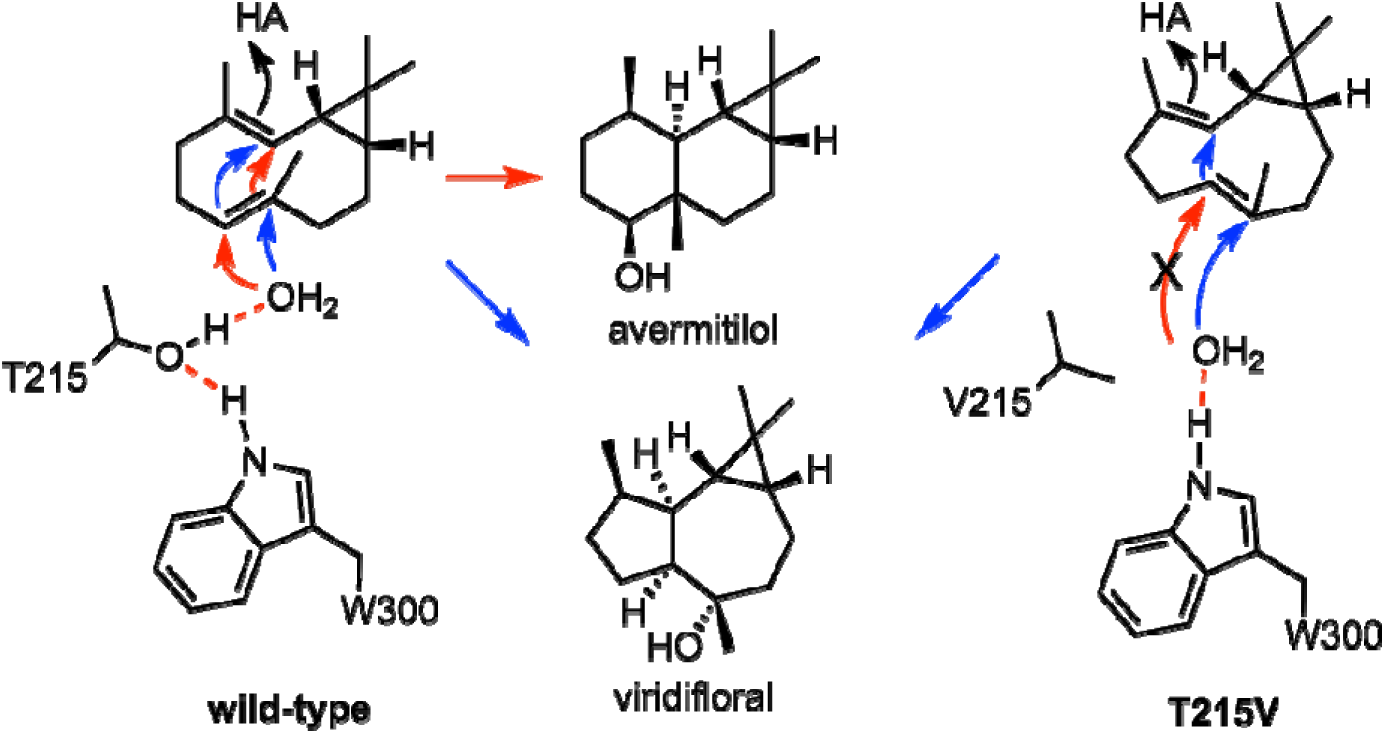
Water management strategies in wild-type SaAS and T215V SaAS. The reactive water molecule indicated likely corresponds to water molecular W1 trapped in the active site (Figures 4C and 5).

Finally, it is interesting that a wide variety of substrate analogues and ligands can be utilized in cocrystallization studies with terpene cyclases.^41,42,43^ Oftentimes, ammonium or aza-analogues can be utilized as mimics of carbocation intermediates and their binding interactions provide key mechanistic inferences regarding the stabilization of carbocation intermediates in catalysis.^44^ The SaAS complexes presented here involve unusual and unnatural ammonium derivatives with adamantane and tetralin skeletons. That such diverse chemical entities are capable of binding with stable conformations in the SaAS active site may reflect untapped potential for further diversity in ligand binding and catalysis. Our future work will continue to explore and expand these biosynthetic possibilities.

## ASSOCIATED CONTENT

### Accession Codes

Atomic coordinates and crystallographic structure factors have been deposited in the Protein Data Bank (www.rcsb.org) with accession codes as follows: SaAS-Mg^2+^_3_-PP_i_-BTAC complex, 9Y6I; T215V SaAS-Mg^2+^_3_-PP_i_-BTAC complex, 9Y6K; A177S SaAS-Mg^2+^_3_-PP_i_–BTAC complex, 9Y6L; SaAS-Mg^2+^_3_-PP_i_-TMAA complex, 9Y6M; SaAS-Mg^2+^_3_-PP_i_-DMTN complex, 9Y6J.

## AUTHOR INFORMATION

## ACKNOWLEDGMENTS

This work is based on research conducted at beamline 17-ID-2 (FMX) of the National Synchrotron Light Source II, a DOE Office of Science User Facility operated for the DOE Office of Science by Brookhaven National Laboratory under Contract DE-SC0012704. The Center for BioMolecular Structure (CBMS) is primarily supported by the National Institutes of Health, NIGMS, through a Center Core P30 Grant (P30GM133893) and by the DOE Office of Biological and Environmental Research (KP1605010).

## References

1. Whitehead, J. N., Leferink, N. G. H., Johannissen, L. O., Hay, S., Scrutton, N. S. (2023) Decoding catalysis by terpene synthases. ACS Catal. 13, 12774–12802.

2. Christianson, D. W. (2017) Structural and chemical biology of terpenoid cyclases. Chem. Rev. 117, 11570–11648

3. Christianson, D. W. (2006) Structural biology and chemistry of the terpenoid cyclases. Chem. Rev. 106, 3412–3442.

4. Tu, Y. (2016) Artemisinin – a gift from traditional Chinese medicine to the world (Nobel lecture). Angew. Chem. Int. Ed. Engl. 55, 10210–10226.

5. Peralta-Yahya, P. P., Ouellet, M., Chan, R., Mukhopadhyay, A., Keasling, J.D., Lee, T. S. (2011) Identification and microbial production of a terpene-based advanced biofuel. Nature Commun. 2, 483.

6. McAllister, J. C., Adams, M. F. (2010) Mode of action for natural products isolated from essential oils of two trees is different from available mosquito adulticides. J. Med. Èntomol. 47, 1123–1126.

7. Cane, D. E. (1990) Enzymic formation of sesquiterpenes. Chem. Rev. 90, 1089–1103.

8. Miller, D. J., Allemann, R. K. (2011) Sesquiterpene synthases: passive catalysts or active players? Nat. Prod. Rep. 29, 60–71.

9. Aaron, J. A., Christianson, D. W.(2010) Trinuclear metal clusters in catalysis by terpenoid synthases. Pure Appl. Chem. 82, 1585–1597.

10. Starks, C. M., Back, K., Chappell, J., Noel, J. P. (1997) Structural basis for cyclic terpene biosynthesis by tobacco 5-*epi*-aristolochene synthase. Science 277, 1815–1820.

11. Aaron, J. A., Lin, X., Cane, D. E., Christianson, D. W. (2010) Structure of epi-isozizaene synthase from Streptomyces coelicolor A3(2), a platform for new terpenoid cyclization templates Biochemistry 49, 1787–1797.

12. Lesburg, C. A., Zhai, G., Cane, D. E., Christianson, D. W. (1997) Crystal structure of pentalenene synthase: mechanistic insights on terpenoid cyclization reactions in biology. Science 277, 1820–1824.

13. Matos, J. O., Kumar, R. P., Ma, A. C., Patterson, M., Krauss, I. J., Oprian, D. D. (2020) Mechanism underlying anti-Markovnikov addition in the reaction of pentalenene synthase. Biochemistry 59, 3271–3283.

14. Kumar, R. P., Matos, J. O., Black, B. Y., Ellenburg, W. H., Chen, J., Patterson, M., Gehtman, J. A., Theobald, D. L., Krauss, I. J., Oprian, D. D. (2024) Crystal structure of caryolan-1-ol synthase, a sesquiterpene synthase catalyzing an initial anti-Markovnikov cyclization reaction. Biochemistry 63, 2904–2915.

15. Wang, T., Yang, Y., He, M., Liu, M., Huang, J. W., Min, J., Chen, C. C., Liu, Y., Zhang, L., Guo, R. T. (2022) Structural insights into the cyclization of unusual brasilane-type sesquiterpenes. Int. J. Biol. Macromol. 209, 1784–1791.

16. Tantillo, D. J. (2010) The carbocation continuum in terpene biosynthesis – where are the secondary cations? Chem. Soc. Rev. 39, 2847–2854.

17. Cane, D. E., Tillman, A. M. (1983) Pentalenene biosynthesis and the enzymatic cyclization of farnesyl pyrophosphate. J. Am. Chem. Soc. 105, 122–124.

18. Cane, D. E., Pargellis, C. (1987) Partial purification and characterization of pentalenene synthase. Arch. Biochem. Biophys. 254, 421–429.

19. Seemann, M., Zhai, G., de Kraker, J.-W., Paschall, C. M., Christianson, D. W., Cane, D. E. (2002) Pentalenene synthase. Analysis of active site residues by site-directed mutagenesis. J. Am. Chem. Soc. 124, 7681–7689.

20. Zu, L., Xu, M., Lodewyk, M. W., Cane, D. E., Peters, R. J., Tantillo, D. J. (2012) Effect of isotopically sensitive branching on product distribution for pentalenene synthase: support for a mechanism predicted by quantum chemistry. J. Am. Chem. Soc. 134, 11369–11371.

21. Lou, T., Li, A., Xu, H., Pan, J., Xing, B., Wu, R., Dickschat, J. S., Yang, D., Ma, M. (2023) Structural insights into three sesquiterpene synthases for the biosynthesis of tricyclic sesquiterpenes and chemical space expansion by structure-based mutagenesis. J. Am. Chem. Soc. 145, 8474–8485.

22. Blank, P. N., Pemberton, T. A., Chow, J. Y., Poulter, C. D., Christianson, D. W. (2018) Crystal structure of cucumene synthase, a terpenoid cyclase that generates a linear triquinane sesquiterpene. Biochemistry 57, 6326–6335.

23. Chou, W. K. W., Fanizza, I., Uchiyama, T., Komatsu, M., Ikeda, H., Cane, D. E. (2010) Genome mining in *Streptomyces avermitilis*: cloning and characterization of SAV_76, the synthase for a new sesquiterpene, avermitilol. J. Am. Chem. Soc. 132, 8850–8851.

24. Hong, Y. J., Tantillo, D. J. (2011) How many secondary carbocations are involved in the biosynthesis of avermitilol? Org. Lett. 13, 1294–1297.

25. Kabsch, W. XDS. (2010) Acta Crystallogr., Sect. D: Biol. Crystallogr. 66, 125–132.

26. Winter, G., McAuley, K. E. (2011) Automated data collection for macromolecular crystallography. Methods 55, 81–93.

27. Tickle, I. J., Flensburg, C., Keller, P., Paciorek, W., Sharff, A., Vonrhein, C., Bricogne, G. (2016) STARANISO. Global Phasing Ltd.: Cambridge, United Kingdom. https://staraniso.globalphasing.org/cgi-bin/staraniso.cgi.

28. Vonrhein, C., Flensburg, C., Keller, P., Sharff, A., Smart, O., Paciorek, W., Womack, T., Bricogne, G. (2011) Data processing and analysis with the autoPROC toolbox. *Acta Crystallogr.*, Sect. D: Biol. Crystallogr. 67, 293–302.

29. Adams, P. D., Afonine, P. V., Bunkóczi, G., Chen, V. B., Davis, I. W., Echols, N., Headd, J. J. Hung, L.-W., Kapral, G. J., Grosse-Kunstleve, R. W., McCoy, A. J., Moriarty, N. W., Oeffner, R.; Read, R. J., Richardson, D. C., Richardson, J. S., Terwilliger, T. C., Zwart, P. H. (2010) PHENIX: a comprehensive Python-based system for macromolecular structure solution. *Acta Crystallogr.*, Sect. D: Biol. Crystallogr. 66, 213–221.

30. Abramson, J. et al. (2024) Accurate structure prediction of biomolecular interactions with AlphaFold 3. Nature 630, 493–500.

31. Emsley, P., Lohkamp, B., Scott, W. G., Cowtan, K. (2010) Features and development of Coot. *Acta Crystallogr.*, Sect. D: Biol. Crystallogr. 66, 486–501.

32. Chen, V. B., Arendall, W. B., III; Headd, J. J., Keedy, D. A., Immormino, R. M., Kapral, G. J., Murray, L. W., Richardson, J. S., Richardson, D. C. (2010) MolProbity: all-atom structure validation for macromolecular crystallography. *Acta Crystallogr.*, Sect. D: Biol. Crystallogr. 66, 12–21.

33. Gaudreault, F., Morency, L. P., Najmanovich, R. J. (2015) NRGsuite: a PyMOL plugin to perform docking simulations in real time using FlexAID. Bioinformatics 2015, 3856–3858.

34. Lin, X., Hopson, R., Cane, D. E. (2006) Genome mining in *Streptomyces coelicolor*:L molecular cloning and characterization of a new sesquiterpene synthase. J. Am. Chem. Soc. 128, 6022–6023.

35. Krissinel, E., Henrick, K. (2007) Inference of macromolecular assemblies from crystalline state. J. Mol. Biol. 372, 774–797.

36. Requena, V. G., Srivastava, P. L., Miller, D. J., Allemann, R. K. (2024) Single point mutation abolishes water capture in germacradien-4-ol synthase *ChemBioChem 25*, e202400290.

37. Gaynes, M. N., Osika, K. R., and Christianson, D. W. (2024) Structure and function of sabinene synthase, a monoterpene cyclase that generates a highly strained [3.1.0] bicyclic product. Biochemistry 63, 3147–3159.

38. Jung, Y., Mitsuhashi, T., Kikuchi, T., Fujita, M. (2024) Functional plasticity of a viral terpene synthase, OILTS, that shows non-specific metal cofactor binding and metal-dependent biosynthesis. Chem. Eur. J. 30, e202304317.

39. Vedula, L. S., Jiang, J., Zakharian, T., Cane, D. E., Christianson, D. W. (2008) Structural and mechanistic analysis of trichodiene synthase using site-directed mutagenesis: Probing the catalytic function of tyrosine-295 and the asparagine-225/serine-229/glutamate-233 motif. Arch. Biochem. Biophys. 469, 184–194.

40. Li, Z., Zhang, L., Xu, K., Jiang, Y., Du, J., Zhang, X., Meng, L-H., Wu, Q., Du, L., Li, X., Hu, Y., Xie, Z., Jiang, X., Tang, Y-J., Wu, R., Guo, R-T., Li, S. (2023) Molecular insights into the catalytic promiscuity of a bacterial diterpene synthase. Nat. Commun. 14, 4001.

41. Hyatt, D. C., Youn, B.; Zhao, Y., Santhamma, B., Coates, R. M., Croteau, R. (2007) Structure of limonene synthase, a simple model for terpenoid cyclase catalysis. Proc. Natl. Acad. Sci. USA 104, 5360–5365.

42. Ronnebaum, T. A., Gardner, S. M., Christianson, D. W. (2020) An aromatic cluster in the active site of *epi*-isozizaene synthase is an electrostatic toggle for divergent terpene cyclization pathways. Biochemistry 59, 4744–4754.

43. Osika, K. R.; Gaynes, M. N.; Christianson, D. W. (2025) Crystal structure and catalytic mechanism of drimenol synthase, an unusual bifunctional terpene cyclase–phosphatase. Proc. Natl. Acad. Sci. USA 122, e2506584122.

44. Chen, M., Al-Lami, H. S., Gan, L., Niu, B., Hill, C. C., Christianson, D. W. (2013) Mechanistic insights from the binding of substrate and carbocation intermediate analogues to aristolochene synthase. Biochemistry 52, 5441–5452.

